# DNA nanopores as artificial membrane channels for origami-based bioelectronics

**DOI:** 10.1101/2023.03.01.530356

**Authors:** Le Luo, Swathi Manda, Yunjeong Park, Busra Demir, Jesse Vicente, M.P. Anantram, Ersin Emre Oren, Ashwin Gopinath, Marco Rolandi

## Abstract

Biological membrane channels mediate information exchange between cells and facilitate molecular recognition^1-4^. While tuning the shape and function of membrane channels for precision molecular sensing via de-novo routes is complex, an even more significant challenge is interfacing membrane channels with electronic devices for signal readout^5-8^. This challenge at the biotic-abiotic interface results in low efficiency of information transfer - one of the major barriers to the continued development of high-performance bioelectronic devices^9^. To this end, we integrate membrane spanning DNA nanopores with bioprotonic contacts to create programmable, modular, and efficient artificial ion-channel interfaces that resolve the ‘iono-electronic’ disparity between the biotic environment and electronics. Through simulations and experiments, we show that cholesterol modified DNA nanopores spontaneously and with remarkable affinity span the lipid bilayer formed over the planar bio-protonic electrode surface and mediate proton transport across the bilayer. Using the ability to easily modify DNA nanostructures, we illustrate that this bioelectronic device can be programmed for electronic recognition of biomolecular signals such as presence of Streptavidin, without disrupting the native environment of the biomolecule. We anticipate this robust biotic-abiotic interface will allow facile electronic measurement of inter-cellular ionic communication and also open the door for active control of cell behavior through externally controlled selective gating of the channels.

## Main

In biological systems, communication between cells occurs via membrane proteins and ion channels that act as size-selective filters or stimulus-responsive molecular valves to either passively allow or actively control the flow of ions across the cell membrane^10^. Cellular communication often surpasses information processing in electronic devices in efficiency, regulation, and specificity^11,12^. Augmenting electronic devices with biological components can enable one to access, analyze, and respond to intercellular information via data transduction and signal transmission^10,13^. Examples include metal oxide semiconductors integrated with ATPase^14^, carbon nanotubes^15,16^ and silicon nanowires to sense pH^17^, 2D transistors functionalized with gramicidin^18^, organic electrochemical devices with membrane channels^19,20^, and H^+^ selective bioprotonic devices integrated with gramicidin^21^, alamethicin^21^ and light sensitive rhodopsins^22,23^. Synthetic membrane channels can further increase the functionality of these devices with well-defined geometries, durability, robustness, and ease of modification^24,25^. Self-assembled synthetic membrane channels are particularly attractive due to their ease of fabrication^25^. To this end, Watson-Crick pairing based hybridization of single-stranded DNA (ssDNA) can rationally and in a bottom-up manner design self-assembled DNA origami structures^26^ that mimic membrane proteins with sophisticated architectures^27-30^ and varied functionalities^31-33^. Here, we merge synthetic self-assembled DNA nanopores based ion-channels with H^+^ selective Pd-based bioprotonic contacts to create a biotic-abiotic device that records and modulates H^+^ currents traversing across the bilayer membrane (Fig. 1). With the unique programmability of the DNA nanopores, we demonstrate the adaptability of this device for sensing specific biomolecules in their native state via distinctive electronic signals.

**Fig. 1.**
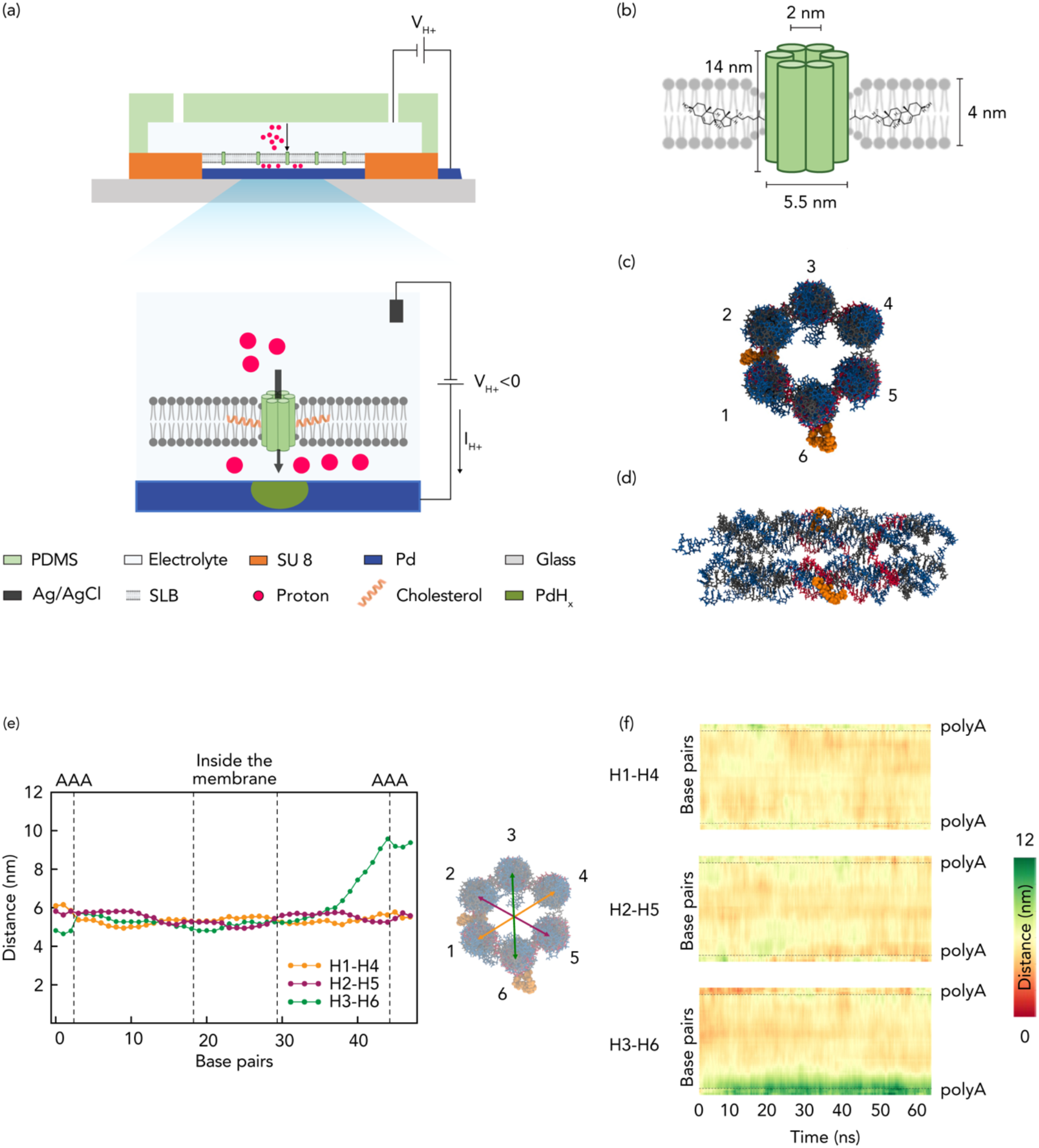
Schematics of bioprotonic devices. (a) Schematic depiction of the bioprotonic device. (b) Schematic representation of a DNA nanopore comprising six-helix bundles and 2-cholesterol anchors. (c) Top view of DNA nanopores with positioned cholesterol anchors. (d) Lateral view of DNA nanopores with positioned base pairs. (e) Simulation of the average distance between the diametrically opposite strands across the length of the nanopore. Yellow for distance between strands 1 and 4, green for distance between strands 2 and 5 and magenta for distance between strands 3 and 6. (f) Average distance heatmap for the pairs indicated in c and e.

### DNA Nanopore Bioprotonics

The DNA nanopore bioprotonic device comprises DNA nanopore ion channels spanning a supported lipid bilayer membrane (SLB) atop a Palladium (Pd) contact integrated with a microfluidic architecture (Fig. 1a and Supplementary Fig.1). A voltage (V_H_^+^) between the Pd contact and Ag/AgCl reference positioned in the solution causes a current of H^+^ between the Pd contact and the solution depending on polarity^34,35^. As previously described, this flow of H^+^ induces the electrochemical formation or dissolution of PdH_x_ that results in a measurable electronic current (I_H+_) in the electronic circuit^21-23^. To create biomimicking ion channels that enable H^+^ transfer across the SLBs, we formed 14 nm long barrel shaped DNA origami nanopores via bottom-up rational design and directed self-assembly (Fig. 1b). To design the DNA nanopores, we first specify the desired 3D geometric shape that closely mimics membrane proteins and stably interfaces with the bilayer membrane^36-38^. We fill the shape from top to bottom with an even number of parallel double helices, held together by periodic crossovers of the strands. The sequences are randomly generated and then rationally down selected to maximize primary interactions as designed and minimize secondary and tertiary complex formations (Supplementary Table 1). The resulting 13 ssDNA aptamers are mixed in equimolar amounts to enable one-pot self-assembly into 6 inter-linked Helix Bundles (6HB) that form the walls the nanopore (Fig. 1b, c, d). We functionalized two of the aptamers with Tetra Ethyl Glycol Cholesterol (TEG-Chol) to provide an anchor for insertion of the hydrophilic DNA nanopores into the hydrophobic environment of the SLB (Fig. 1b, c, d and Supplementary Fig. 2 and 3). Next, we conducted transmission electron microscopy (Supplementary Fig. 2c), dynamic light scattering (Supplementary Fig. 3c) analysis as well as molecular dynamics (MD) simulations to investigate the dimensions and the stability of the 6HB inside the SLB and its pore size under dynamic environments (Fig. 1e and f). The average distance between the diametrically opposite DNA helices across the length of the nanopore is analyzed, as depicted in Fig. 1e, providing insights into the pore size. Additionally, Fig. 1f illustrates the dynamic behavior of these distances, as they change with time on a base pair level within the DNA nanopore. Our analysis reveals that inside the membrane, the center-to-center distances of the opposite helices ranges between 5 and 6 nm. Given that the radius of a DNA double helix is 1 nm, this indicates that the average pore size fluctuates between 3 and 4 nm. Outside the membrane, the DNA helices exhibit increased mobility, resulting in some helices moving apart from one another (Figure 1 e green plot). However, as seen in the Fig. 1f, this phenomenon does not impact the stability of the pore as the TEG-Chol anchors stabilizes the DNA origami inside the SLB. Therefore, we anticipate that the length of the nanopore provides sufficient area for decoration with hydrophobic anchors to enable spontaneous insertion while projecting further beyond the SLB to enable desired interactions at the lip of the nanopore without disrupting its stability within the bilayer. The small inner lumen size facilitates proton transport across the channel while obstructing proteins and other larger biomolecules to remain on the cis (negatively charged) side of the nanopore^36^.

### Control of H^+^ flow with DNA Nanopore Bioprotonics

To validate the DNA nanopore is indeed a H^+^ conductor, we measured the dependence of I_H+_ to V_H+_ in the DNA bioelectronic device (Fig 2a). First, we verified that the bare Pd contact transfers H^+^ at the solution interface (Fig 2a-i). To do so, we recorded I_H+_ as a function of V_H+_ with the following sequence as previously described^21^. In the first step, V_H+_= - 400 mV for 600 seconds so that H^+^ flow from the solution into the Pd contact to form PdH_x_ (Fig. 2a-i) as indicated by I_H+_= - 131 ± 26 nA (Fig. 2b). In the second step, V_H+_ to 0 mV to transfer H^+^ from the PdH_x_ contact into the solution. Here, I_H+_ indicates the prior formation of PdH_x_ that allows H^+^ to transfer from the surface back into the solution even at V_H+_= 0 mV because at a neutral pH the protochemical potential of H^+^ in the PdH_x_ contact is higher than the protochemical potential of H^+^ in the solution^39^. Second, we confirmed that the SLBs create the barriers and block H^+^ transport from the solution into the Pd surface to make sure that when we insert the DNA nanopore we measured H^+^ transport across the nanopore instead of SLBs (Fig. 2a-ii), as indicated by I_H+_= -15 ± 8 nA (Fig. 2b). The measured current, referred to as the leakage current, indicates that few H^+^ diffuse and leak across the bilayer membrane, possibly through the surface defects and are reduced at the Pd surface. We confirmed the characteristics of the Pd contact with Electrochemical Impedance Spectroscopy (Supplementary Fig. 4). After addition of DNA nanopores modified with 2 cholesterol handles (6HB-2C) in the solution, we expect the DNA nanopores spontaneously insert into the lipid bilayer (Fig 2a-iii). This insertion results in I_H+_=-44 ± 8 nA for V_H+_ = -400 mV (Fig. 2b), which is 3 times larger than I_H+_ for the SLBs coated Pd indicating that the DNA nanopores provide a pathway for H^+^ to move across the SLB. For all measurements, to avoid the accumulation of protons on Pd contact, in the second sequence, we set the V_H+_ to 0 mV. The higher photochemical potential than the electrolyte led to the release of protons into the electrolyte and with positive I_H+_. As predicted, DNA nanopores without any cholesterol handles (Fig. 2a-iv) did not insert into the SLB corroborated by the same I_H+_ as recorded for the naked SLB (Fig. 2c). Nanopores with one or three cholesterol handles (6HB-1C, 6HB-3C) (Fig. 2a-v, vi) also did not insert into the SLB (Fig. 2d). It is likely that 6HB-1C does not insert into the SLB because one cholesterol handle is not enough to drive the hydrophilic DNA nanopore into the hydrophobic SLB^40^. However, with the same reasoning one would we expected to see even better insertion for 6HB-3C compared to 6HB-2C. It is likely that the increased hydrophobicity of 6HB-3C drives its aggregation in solution to minimize interaction with water and makes it unavailable for insertion into the SLB. This aggregation is confirmed by multiple bands 6HB-3C in gel electrophoresis (Supplementary Fig. 4a) and a hydrodynamic radius eight times larger for 6HB-3C compared to 6HB as measured by DLS (Supplementary Fig 4c).

**Fig. 2.**
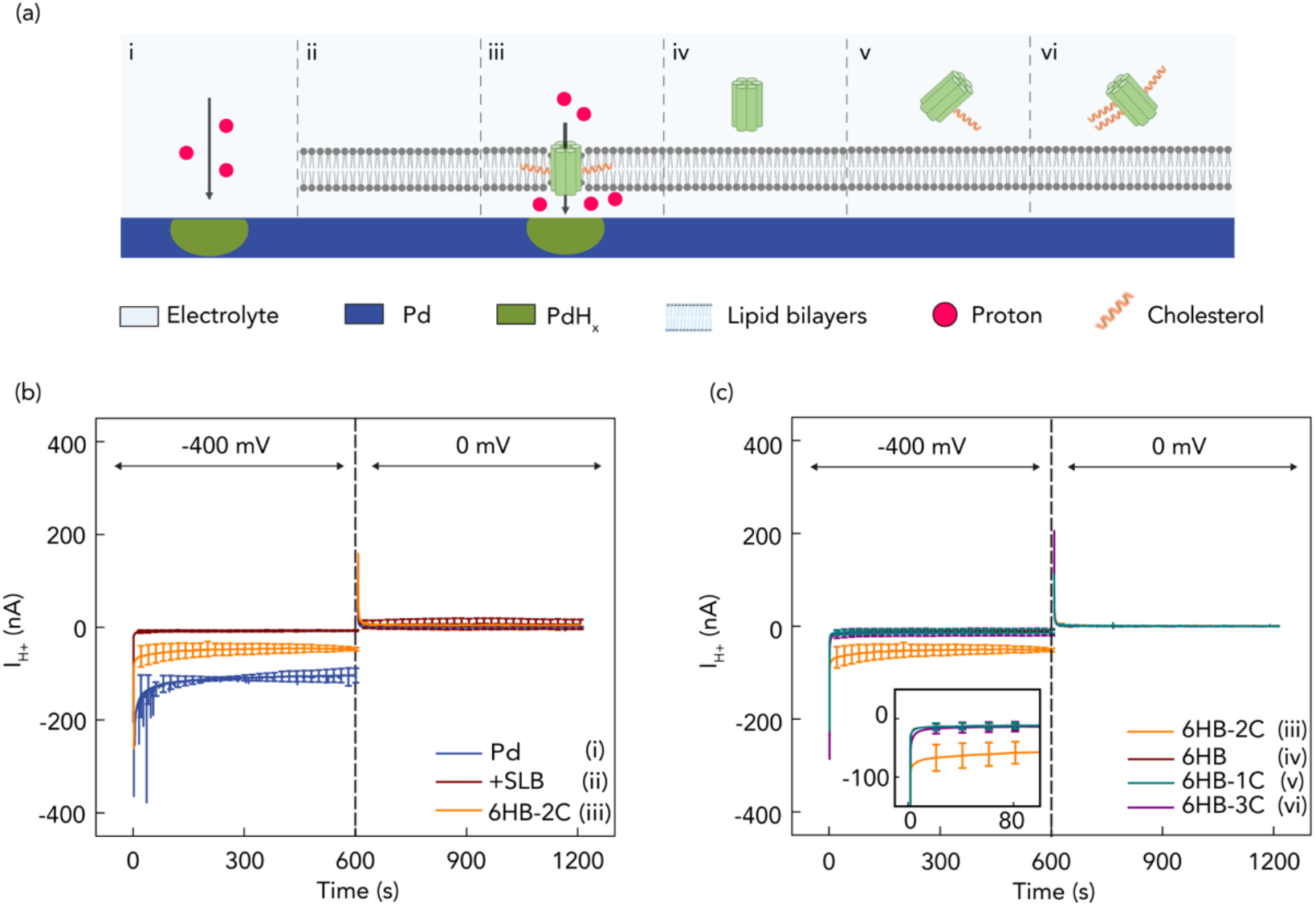
Schematics of control of H^+^ by membrane spanning DNA nanopores. (a) i) Pd contact with electrolyte solution; ii) Pd contact coated with SLBs; iii) DNA nanopores with 2-cholesterol anchors (6HB-2C); iv) DNA nanopores without cholesterol anchor (6HB); v) DNA nanopores with 1-cholesterol anchor (6HB-1C); vi) DNA nanopores with 3-cholesterol anchor (6HB-3C). (b) I_H+_ versus time plot for V = -400 mV and V = 0 mV. Blue trace Pd, red trace SLB, Orange trace 6HB-2C. The I_H+_ = -131 ± 26 nA with bare Pd decreased to - 15 ± 8 nA with SLBs that indicates formed bilayers inhibit H^+^ transfer from the bulk solution to the Pd/solution interface. The I_H+_ = -44 ± 8 nA with 6HB-2C confirmed that the nanopore channels support the H^+^ transport. (c) I_H+_ versus time plot in different situations of Fig.2a under V = -400 mV and V = 0 mV. Orange trace 6HB-2C (2a-iii), red trace 6HB (2a-iv), cyan trace 6HB-1C (2a-v), and purple trace 6HB-3C (2a-vi). Under -400 mV, we also measured I_H+_ = -11 ± 2 nA, -16 ± 3 nA, and -11 ± 3 nA with 6HB, 6HB-1C and 6HB-3C, respectively. Only 6HB-2C provides created pathway to facilitate the flow of H^+^ to the Pd/solution interface. For all measurements, we switched the voltage to 0 mV, roughly after 600 s from the first instance of measurement, the H absorbed in Pd is oxidized to H^+^ and released back into the solution, allowing the current measured to return to 0 nA.

### Programming the DNA nanopores for biomolecular sensing

DNA self-assembly allows to program any desired functionality in the DNA nanopores by designing ad-hoc DNA sequences. As proof-of-concept biomolecular sensing, we program DNA nanopores for the detection of Streptavidin (S-avidin) by including a biotin functionality ^41^. We did so by functionalizing 6HB-2C nanopores using ssDNAs modified with a Biotin tag at their 5’ ends followed by DNA hybridization to obtain the formation of 6HB-2C-2B nanopores with a biotin tag at either end of the nano barrel (Fig. 3a-i). As expected, 6HB-2C-2B nanopores insert themselves into the SLB and result in a large I_H+_= -92±15 nA for V_H+_= -400 mV indicating that the nanobarrel inside the DNA nanopore aids H^+^ transport across the SLB (Fig 3aii-b). However, with S-avidin in solution the binding event of the 5 nm S-avidin with the biotin handle^42^ on the DNA nanopore effectively occludes the nanobarrel impeding H^+^ transport across the SLB as indicated by a reduction of I_H+_ to the level of SLB leakage I_H+_. As control, exposing non biotinilated 6HB-2C to the same S-avidin concentration in solution does not cause any appreciable change in I_H+_ (Fig. 3a-iii, b) because the S-avidin has no binding site available on the DNA nanopore that would result in the occlusion of the nanobarrel. While this is a simple proof-of-concept, by leveraging the programmability offered by the DNA nanostructures, we engineered the nanopores to demonstrate an electronic sensing response to a specific analyte in its native environment without the need for modifying the analyte. With the large variate of available DNA sequencies and potential for adding a variety of multiple functionalities, this approach can be extended to electronically sense multiple analytes simultaneously.

**Fig. 3.**
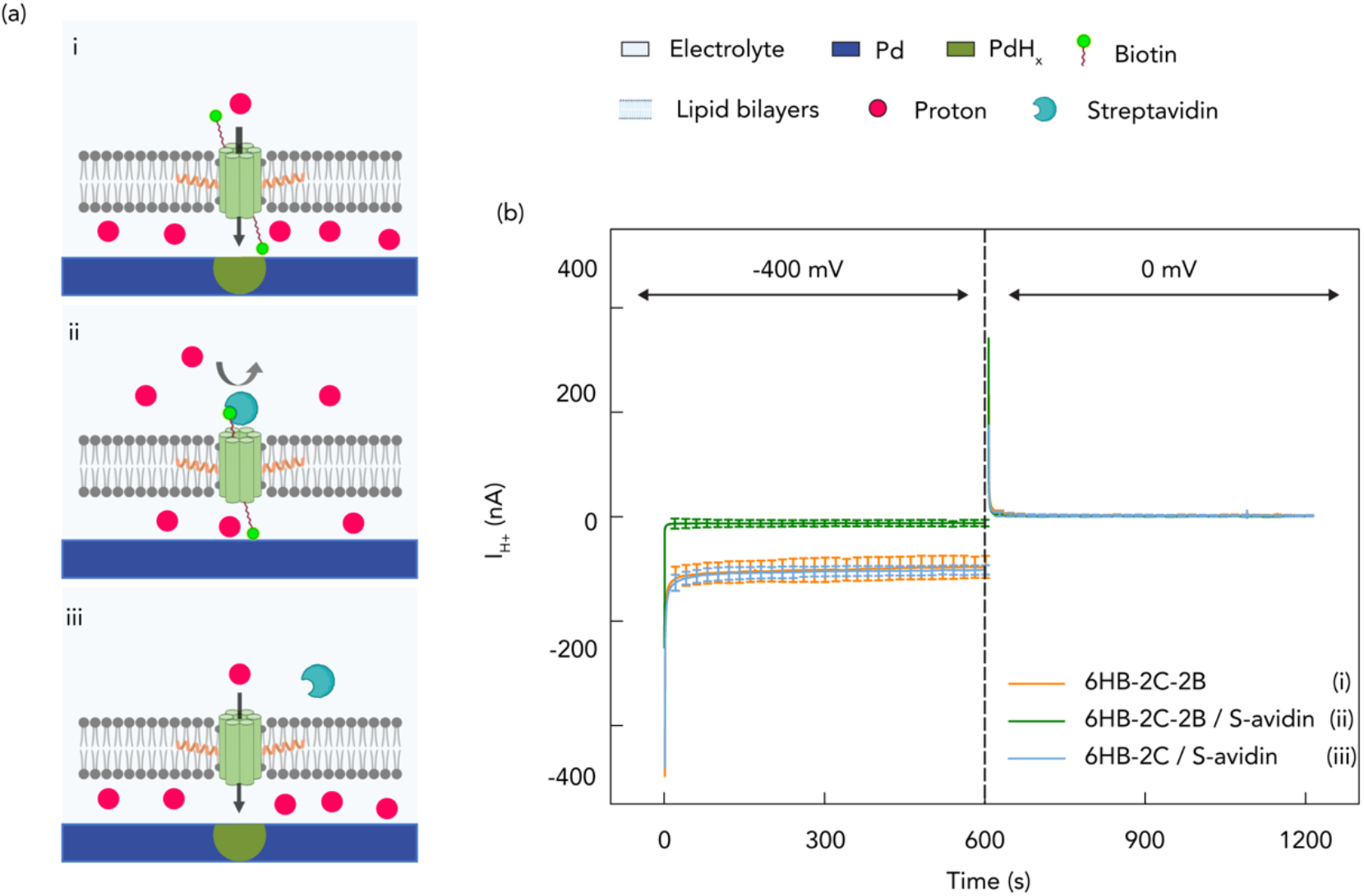
Schematics of bioprotonic devices with biotin-Streptavidin. (a) i) 2-cholesterol handled DNA nanopores with biotin in the absence of streptavidin (6HB-2C-2B). Created pathway facilitates H+ transfer without inhibition of binding due to absence of streptavidin; ii) 2-cholesterol-handled DNA nanopores with binding of biotin-streptavidin (6HB-2C-2B/S-avidin). H^+^ transfer is inhibited by blocked pore channels; iii) 2-cholesterol-handled DNA nanopores without biotin in the presence of streptavidin (6HB-2C/S-avidin). The pores are not blocked by binding due to lacking biotin. (b) I_H+_ versus time plot for V = -400 mV and V = 0 mV. Orange trace 6HB-2C-2B (3a-i), green trace 6HB-2C-2B/S-avidin (3a-ii) and blue trace 6HB-2C/S-avidin (3a-iii). We measured I_H+_ = -92 ± 15 nA, -13 ± 4 nA and -99 ± 2 nA with 6HB-2C-2B, 6HB-2C-2B/S-avidin and 6HB-2C/S-avidin, respectively.

### A model for the rate constant of association and dissociation of DNA nanopores self-insertion

To better understand the dynamics of DNA nanopore insertion with the SLB, we created a model based on Langmuir’s equation and absorption/desorption kinetics^43,44^ to analyze the insertion process of the DNA nanopore in the lipid bilayer. In this model, we describe DNA nanopores in solution (n) and lipid bilayer sites where the nanopores can be absorbed (l) as being initially separate (Eq. 1 left side). Upon insertion of the DNA nanopore into the lipid bilayer, the DNA nanopore and the lipid bilayer sites are conjoined together and we describe this entity as nl (Eq. 1 right side).

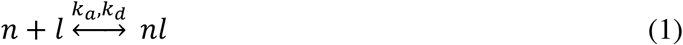

The rate constant k_a_ (M^-1^s^-1^) describes the absorption reaction of the DNA nanopore into the lipid bilayer and the rate constant k_d_ (s^-1^) describes the desorption reaction. From this model, we expect that more DNA nanopores in solution (n) correspond to a higher number of DNA nanopores inserted into the lipid bilayer (nl) resulting in I_H+_ to increase as a function of DNA nanopore concentration (C_n_) (Figure 4a). Given the large number of DNA nanopores compared to the absorption area of the lipid bilayer, we assume C_n_ to be constant throughout the absorption process. To fully understand the absorption and desorption kinetics, we need to now derive k_a_ and k_d_. To do so, we introduce the differential form of the Langmuir equation:

**Fig. 4.**
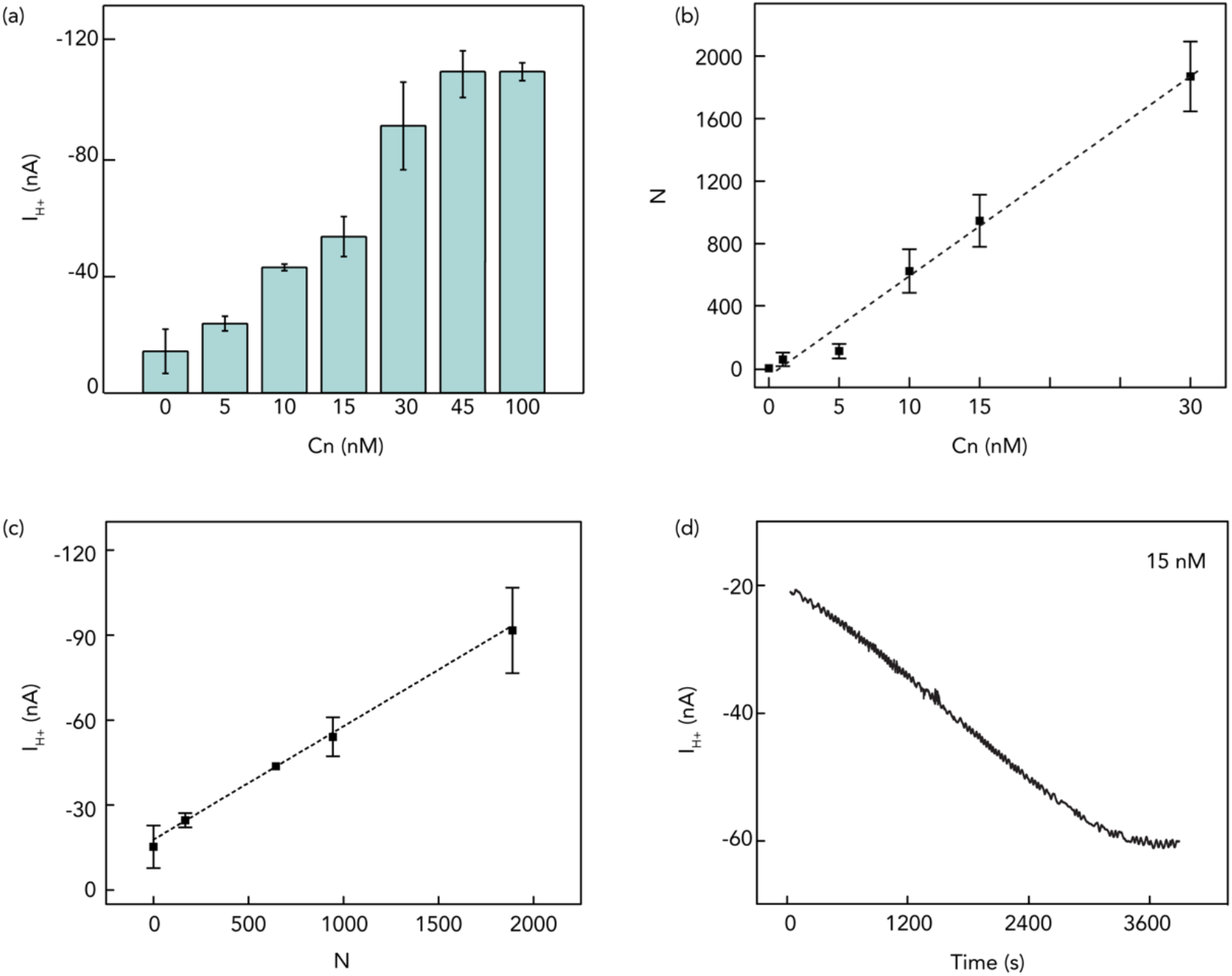
Illustration of DNA nanopores characteristics. (a) I_H+_ versus C_n_ plot (V_H+_ = -400 mV). (b) The number of inserted 6HB-2C nanopores versus the introduced 6HB-2C concentration plot. The value of the slope in the plot is 64. (c) I_H+_ versus Number of inserted 6HB-2C nanopores under - 400 mV. The value of the slope in the plot is 4ξ10^−11^. (d) I_H+_ versus time plot during 15 nM 6HB-2C insertion process under -400 mV.

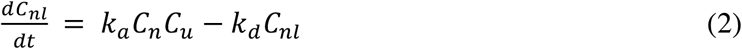

where C_n_ and C_nl_ are the DNA nanopore concentrations in solution and lipid bilayers, respectively, and C_u_ represents the unoccupied site concentration in the SLB. Since C_u_ is an unknown that we are not able to derive experimentally, we write (2) as:

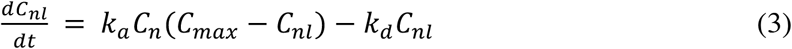

Where C_max_ = *C*_*u*_ + *C*_*nl*_ and C_max_ is the maximum value of C_nl_. We derive C_nl_ by counting the number of inserted DNA nanopores (*N = C*_*nl*_*V*_*l*_*A, V*_*l*_ *= the volume of lipids, A = Avogadro’s number*) as a function of C_n_ at equilibrium using fluorescent microscopy on fluorescently tagged nanopores (Fig. 4b and Supplementary Fig. 5). Unfortunately, we are not able to measure C_max_ using fluorescent microscopy for C_n_ > 30 nM because the inserted DNA nanopores are too close to each other and difficult to count. In Figure 4a, we have shown that I_H+_ increases with increasing C_n_ for C_n_< 45 nM and then I_H+_ plateaus even if we increase C_n_ up to100 nM. We assume that for C_n_> 45 nM, C_nl_= C_max_. To calculate C_max_ from I_H+_, we then model the DNA nanopores as resistors in parallel:

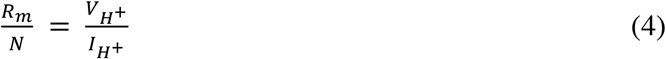

and from equation 4 and the slope of Figure 4c we calculate R_m_ = 1 × 10^10^ Ω and G_m_ = 1/R_m_ = 100 pS, the resistance and conductivity of each individual nanopore. These values are consistent with the conductivity of artificial and natural membrane channels^45,46^. Using the calculated value of R_m,_ N_max_ = 2350 nanopores per device and C_max_=2 nM. To conclude the derivation of k_a_ we then observe experimentally *C*_*nl*_*/dt* by recording I_H+_ as a function of time introducing the DNA nanopores in the solution for t=0 (Fig. 4d). We can thus assume C_nl_=0 and Eq 3 simplifies to:

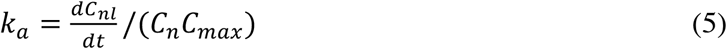

Using *N = C*_*nl*_*V*_*l*_ *A* and equation 3 we can express *dC*_*nl*_*/dt* as:

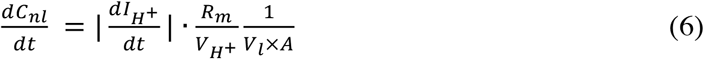

Combing equation 5 and 6 we can express *k*_*a*_ as:

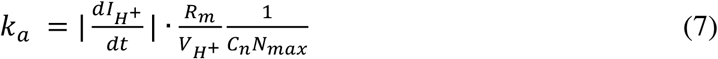

From the slope of Fig 4d at *t = 0*, we calculate *k*_*a*_ *= 8*.*5 × 10*^*3*^ *M*^*-1*^*s*^*-1*^.

We then look at time t when the system reaches dynamic equilibrium and *dC*_*nl*_*/dt= 0* and write:

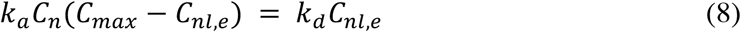

where C_nl,e_ is the adsorbate concentration in bilayers at equilibrium. We derive C_nl,e_ from I_H+_ and calculate k_d_ = 1.9 × 10^−4^ s^-1^. We then calculate the apparent dissociation constant to be *K*_*D*_ *= k*_*d*_*/k*_*a*_ = 22 nM. The apparent dissociation constant indicates a high affinity of the 6HB-2C to the SLBs and is higher than the affinity of most protein-ligand interactions (100 μM — 100 nM)^45,47,48^.

## Discussion

We have successfully demonstrated a programmable bio-protonic device with membrane-spanning DNA nanopore ion channels as molecularly precise interconnects that measure and control the H^+^ transfer across lipid bilayer interface. By carefully tuning the size of the nanopore and modifying its surface, we introduced a new class of molecular signal transductors that can be formed via self-assembling and can self-insert into membranes. For the first time, we explored the kinetics of DNA nanopore experimentally through ensemble experiments. Variations of this next-generation bio-protonic device, such as ligand modified DNA nanopore or external energy triggered conformational pore change mechanism will facilitate high-throughput multi-omics, including measuring DNA, RNA, protein, or small molecule binding. To make the nanopore respond to the presence of a molecule (DNA, RNA, proteins, or small molecule), we can design a DNA or RNA aptamer (or other affinity binders) on the cap of the nanopore. Free aptamer will remain unstructured and does not significantly affect the proton transport through the pore, whereas upon binding its target, it will undergo a significant conformational change and significantly reduced the nominal diameter of the ion channel leading to a marked reduction in the measured proton conductivity. We anticipate this device to be expanded into a multiplexed proteomic and metabolomics chip composed of a precisely decorated array of active DNA nanopore ion channels. Due to the ease of DNA functionalization, each distinct subarray within the array may be programmed to respond to a particular ligand (protein or small molecule) based on the choice of aptamer on it. Because each DNA nanopore electrode in this setting will be gated by a single copy of a known molecule, the concentration of a given ligand in solution can be directly quantified based on the rate at which the ligands gate their corresponding nanopores. With this approach, this device will open pathways for measuring absolute numbers of molecules in a highly multiplexed and scalable setting. Furthermore, this device will enable an ideal abiotic-biotic interface to measure inter and intra-cellular electrical measurements. Specifically, active, or passive DNA nanopore-electrodes can be designed to spontaneously insert into the cultured cells instead of lipid bilayers so that each nanopore creates a precise electrical interface between the cells and the bio-protonic electrode. Unlike patch clamp experiments, or biological nanopore based bioelectronic devices, where electrical current signals from the cells can only be monitored, this device will allow us to control the transport within a cell or between cells through ligand specific gating of the functionalized DNA nanopores. Thus, this device will enable real time stochastic sensing of molecules inside and outside of the cellular environments.

### Materials

1,2-dioleoyl-sn-glycerol-3-phosphocholine (DOPC, Avanti Polar Lipids), 1,2-dipalmitoyl-sn-glycerol-3-phosphoethanolamine-N-(lissamine rhodamine B sulfonyl) (Fluorescent liposomes, Avanti Polar Lipids) were used as received for formation of supported lipid bilayers. Unmodified ssDNA oligos (for sequences see Table S1) in 25 nmole scale with standard purification, 3’ TEG-Chol modified ssDNA oligos (for sequences see Table S1) in 100 nmole scale with HPLC purification and 5’Bn modified ssDNA oligos (for sequences see Table S1) in 25nmole scale with standard purification were obtained from Integrated DNA Technologies. TE buffer 10x (pH = 8.0), MgCl_2_ ∙ 6H_2_O, 3-aminopropyl-triethoxy-silane (APTEs) and PBS (pH = 7.5) were purchased from Sigma-Aldrich. Streptavidin was purchased from Thermo Fisher Scientific. The Ag/AgCl reference electrode (RE) and counter electrode (CE) were from Warner Instruments. Glass wafers, 4-in diameter were obtained from University Wafer Inc.

## Methods

### Device architecture, fabrication and characterization

Bioprotonic devices were fabricated with conventional soft- and photo-lithography on a 100 nm thick layer of glass. The SU-8 insulating channel is 10 μm thick and the PDMS microfluidic channel is 100 μm on each chip. The contact area of Pd contacts is 0.25 mm^2^ (500 × 500 μm) with a thickness of 100 nm for significant interfacing with lipid solution. Pd is deposited on top of 5 nm Cr adhesion layer. The microfluidic channel confines the flow of liquid to the top of the Pd contact and provides space to insert RE and CE (Supplementary Fig. 1). In EIS measurement, Autolab was also used to record impedance spectra in the frequency range between 0.1 Hz–100 kHz. An AC voltage of 0.01 V and a DC voltage of 0 V versus OCP (open circuit potential) were applied (Supplementary Fig. 3).

### SLB formation

DOPC liposomes were extracted and dried from a vial containing DOPC and chloroform via using nitrogen to blow it. And thus, the vial was put into a vacuum chamber for at least 6 hours to dry DOPC extremely. Followingly, PBS buffer solution (pH = 7.5) was added into the vial for rehydration with the exact density (1 mg ml^-1^). Sonication and vortex promote dissolution of DOPC in buffer solution and then 220 nm sterilizing filters purchased from Millex confine the size of vesicles. Before the deposition of SLBs on Pd contacts, the surface was hydrophilized by oxygen plasma^49^. The vesicle solution was introduced and dispensed in the microfluidic channel and the device was gently agitated for at least 8 h in high relative humidity (∼ 95 % RH) to ensure vesicle fusion^50,51^ and the SLB formation, followed by rinsing with buffer solution to wash away vesicle residue that was unfused. Essentially, the SLBs mimic cell membranes, electrically insulate the Pd contact and divides the solution into two volumes: cis and trans. SLBs are not in direct contact with the surface of the solid substrate because of a very thin hydration layer of 1—2 nm thickness between Pd contact and SLBs on this cis-side^52^. The separation offered by this thin layer facilitates the insertion of ion-channels such as the DNA nanopores by supplying lubrication and mobility to the SLBs^53^.

### Fluorescence imaging and fluorescence recovery after photobleaching measurements (FRAP)

The formation and quality of SLBs is validated by Fluorescence Recovery After Photobleaching (FRAP) (Supplementary Fig. 6). Fluorescence imaging and FRAP experiments were performed on confocal microscopy (Leica, SP5 Confocal Microscope) with a 63× water immersion objective. DNA nanopores were tagged with 488 Atto fluorophores. 488 nm Ar laser was used for fluorescence imaging and 543 nm and 594 nm HeNe laser was used for photobleaching. Both samples were flushed with PBS several times for removing excess fluorophore. A 20 μm diameter spot in the supported lipid bilayer was photobleached and its fluorescence intensity recovery was monitored for 30 minutes. The fluorescence intensity of diameter and changes over time were analyzed with image J and fitted using a Gaussian function^54^. The diffusion coefficient was calculated with the below equation:

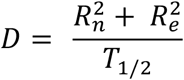

where R_n_ is the nominal radius from the user defined spot, *R*_*e*_ is the effective radius from bleached radius right after the bleaching process, *T*_*1/2*_ is half time to recovery and the diffusion coefficient was 8.52 μm^2^/sec.

### DNA nanopores folding and characterization

The 6HB-2C DNA nanopore was assembled by heating and cooling an equimolar mixture of 11 unmodified and 2 TEG-Chol modified DNA strands (for sequences see Table S1). 10 uL of each of 1 µM ssDNA were mixed along with 6 uL of 200 mM MgCl_2_, 10 uL of 10x TE (pH = 8.0) and MQ water to prepare a 100 uL folding mixture. The mixture was divided into 50 uL aliquots so that the solution maintains an even contact with the heating elements of the thermocycler. They were first heated to a temperature of 95°C and then sequentially cooled to 16 °C by reducing the temperature at a rate of 0.13 °C per minute. For 6HB control nanopores without cholesterol anchors (6HB) and other controls such as 6HB-1C, and 6HB-3C, 6HB-2B, 6HB-2C and fluorescent tag nanopores, the sequences were appropriately modified (for sequences see Supplementary Table S1).

The self-assembled structures were then characterized to confirm the correct and successful formation of the DNA nanopore. Since the structures were formed from equimolar ratios of ssDNA strands, purification was not necessary. The concentration of the resulting double stranded DNA (dsDNA) nanostructures was analyzed with a spectrophotometer using UV absorbance spectra. Native gel electrophoresis was performed to verify the completeness of the folded structure and to verify the migration of the control nanopores without any cholesterol vs. migration of 6HB-1C, 6HB-2C, and 6HB-3C nanopores (Supplementary Fig. 4a). The 6HB-2C nanopores yielded a band, which migrated to the similar height as a control nanopore without any cholesterol anchors (Supplementary Fig. 4a, lanes 3 and 5 respectively, main band migrating at 300 bp marker). Furthermore, dynamic light scattering (DLS) established the monomeric nature of the nanobarrels, as only a single peak with an average hydrodynamic radius of 9.73 nm was observed (blue curve in Supplementary Fig. 4c). For biotin modified nanopores, 6HB-2B and 6HB-2B-2C, the gel electrophoresis showed the accessibility of the biotin tags as slower migration patterns and dimer/quadrate aggregation patterns were observed in presence of excess Streptavidin protein (1x:20x concentration ratios) (Supplementary Fig. 4b lanes 6 and 8). No such migration pattern changes were observed in non-biotinylated nanopores, showing that they were appropriate as controls (Supplementary Fig. 4b lanes 7).

### Simulation

We perform molecular dynamics (MD) simulations with NAMD software^55,56^ using periodic boundary conditions. 6HB DNA origami design is generated with caDNAno and converted into all atom structures using automated conversation program which is available at the nanoHub web site. We covalently bind cholesterol-TEG (chol-TEG) extensions to 3’ends of designed staple strands by using the ‘patches’ provided in the NAMD tutorial^57^. We, then, inserted the chol-TEG conjugated 6HB DNA origami nanopore into the pre-equilibrated DOPC lipid bilayer membrane using the CHARMM-GUI website^58^. CHARMM 36^59^ and CGenFF^60^ force fields were used to define chol-TEG conjugated DNA origami structure. We placed the whole system inside 0.15 KCl electrolyte after removing overlapping lipid and water molecules. For water molecules and ions TIP3P^61^ force field was used. After generating the initial system, we minimized the energy of lipid molecules for 50000 steps by keeping chol-TEG conjugated DNA origami structure fixed. Next, we minimized the energy of the whole system while keeping the chol-TEG conjugated DNA origami harmonically restrained using the exponent of 2 for the harmonic constraint energy function for another 50000 steps. We released all the harmonic constraints and equilibrated the whole system for 3 ns prior MD production runs. Finally, the whole system was simulated for 64 ns at 295 K with a 2 fs timestep by saving the coordinates at every 4 fs. During the simulations, the VDW cutoff value is taken to be 12 Å. Electrostatic interactions are computed using the PME method^62^, and the SHAKE algorithm is applied to keep H bonds rigid.

## Supporting information

Supplemental figure 1 will be used for the link to the file on the preprint site

---

**Supplementary Fig. 1.**
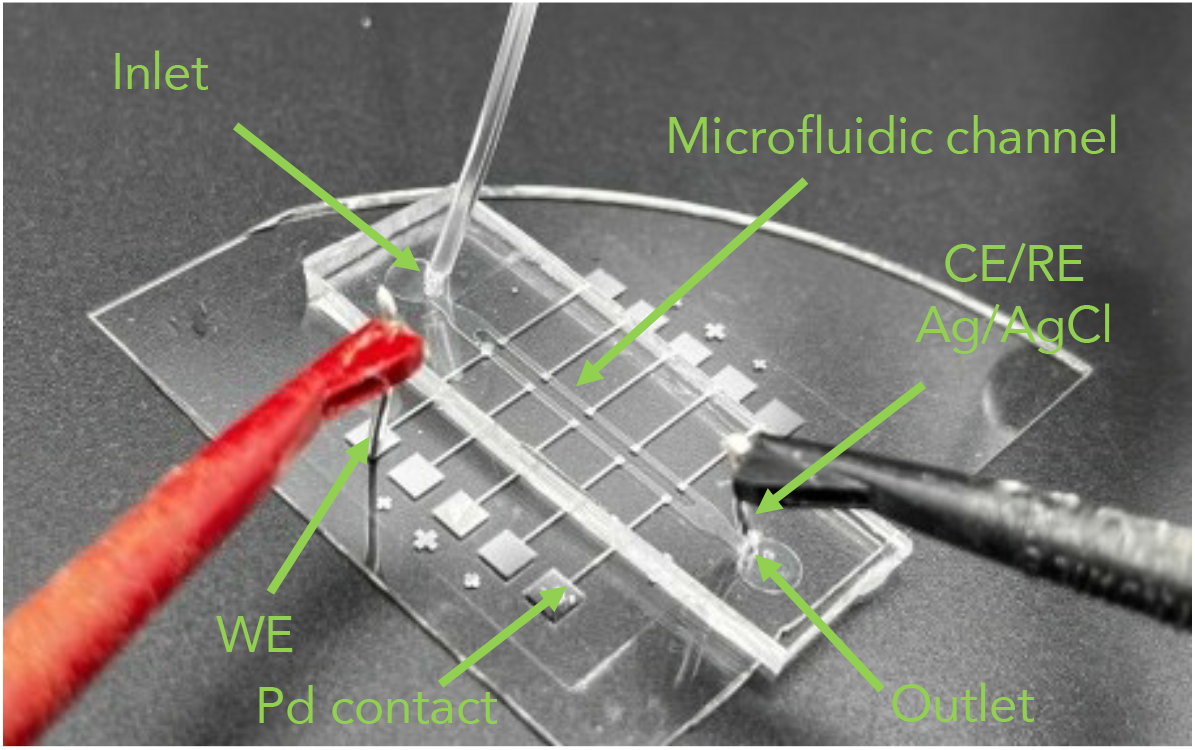
Optical image of the bioprotonic device.

**Supplementary Fig. 2.**
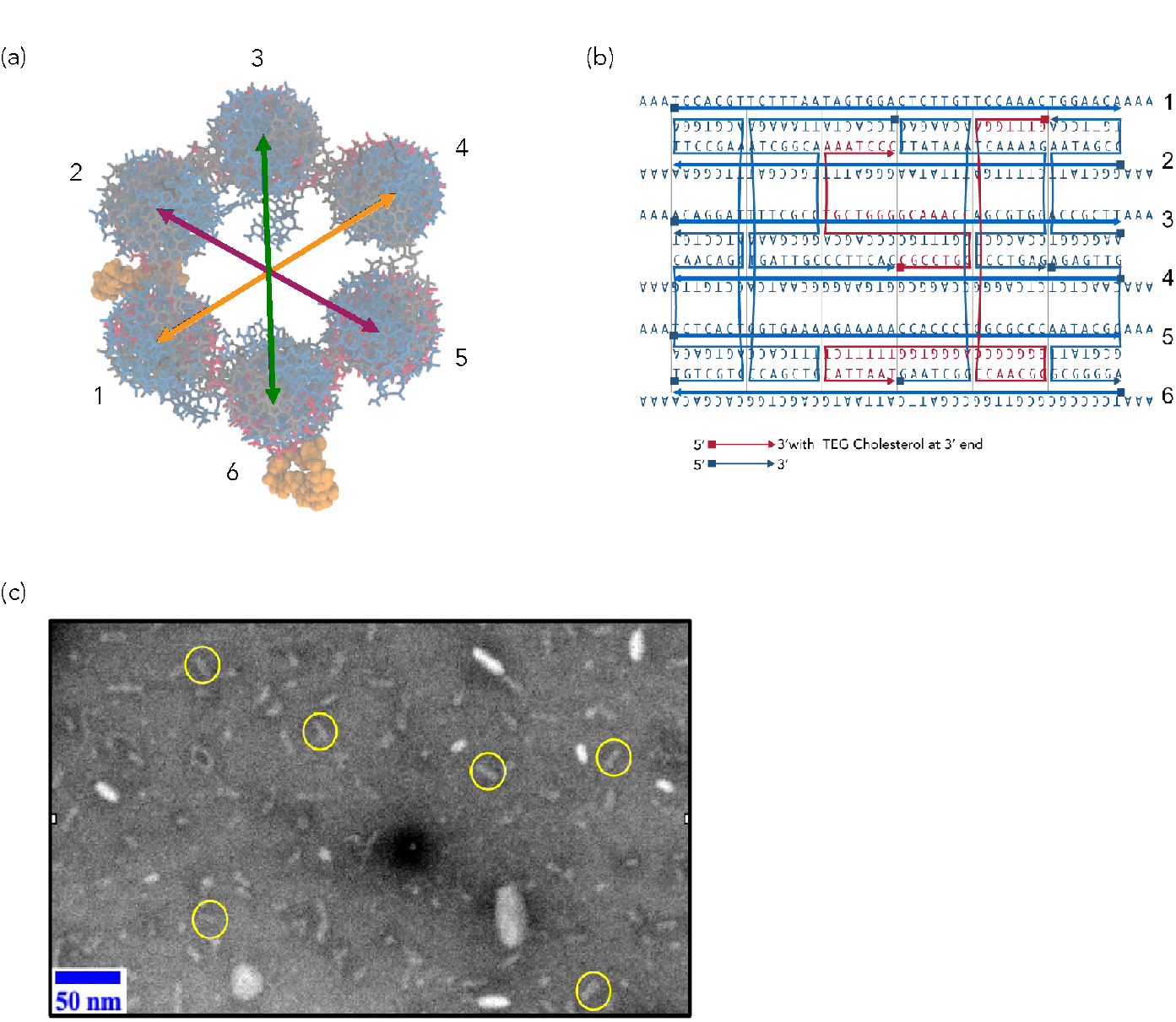
6HB-2C nanopore design, conformation and simulations. (a) Simulation of cholesterol moieties on helices 2 and 6 from an axial view with respect to the six helix bundles. (b) Sequence design and strand crossover details. Red strands indicate those oligos that have been modified at the 3’ end with Tri-ethylene Glycol (TEG) cholesterol moieties. Blues strands indicate oligos without any modifications. Squares indicate the 5’ end while triangles indicate the 3’ end of DNA. (c) Negatively stained TEM micrograph of the 6HB nano-barrels. Yellow circles show the nanopores in a flat orientation.

**Supplementary Fig. 3.**
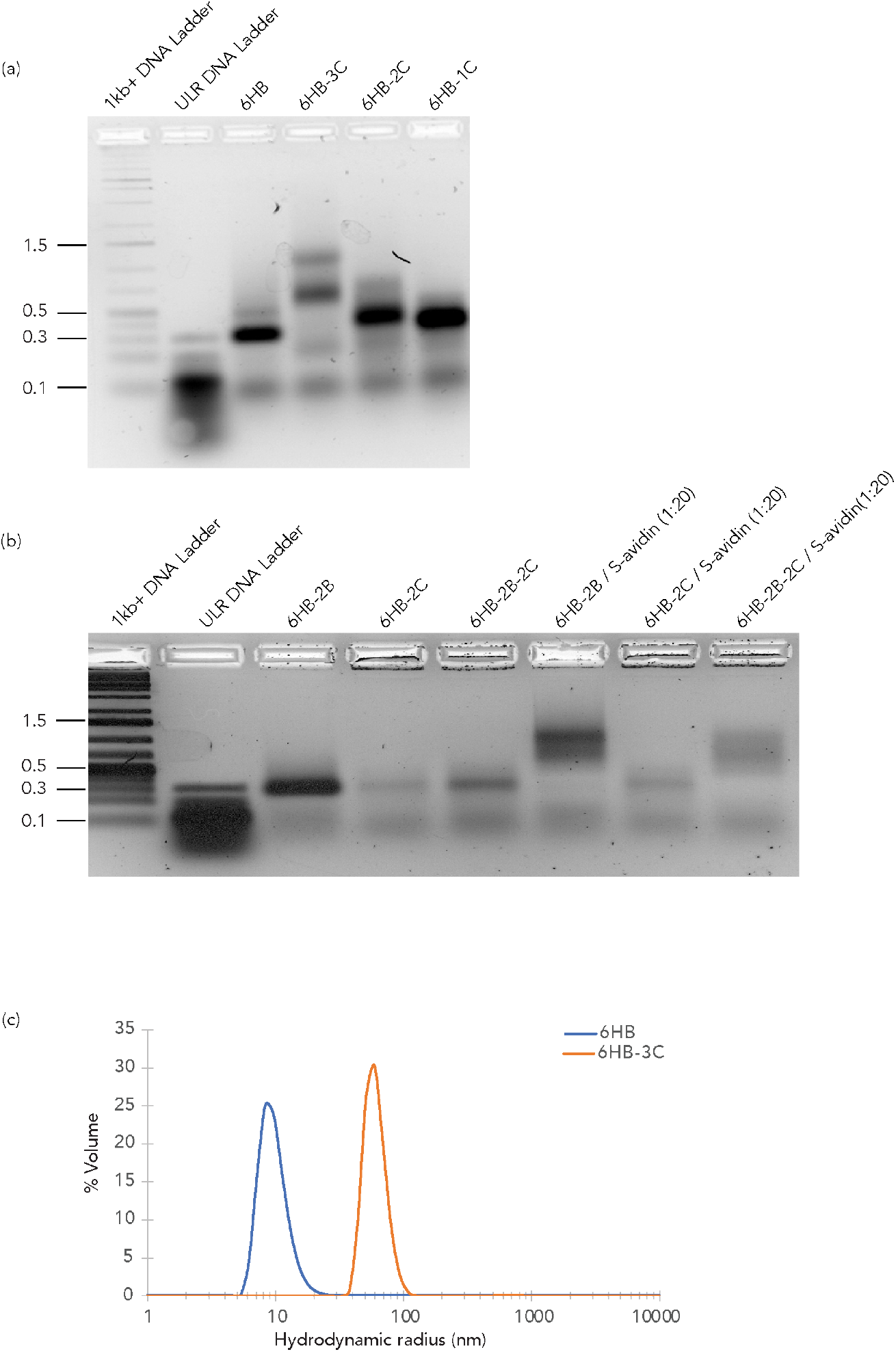
Design and verification of DNA nanopores with cholesterol handles. (a) Electrophoresis characterization (2% Agarose gel) of the nanopores with different number of cholesterol tags around the midsection of the nano-barrel. Lane 1 and 2, DNA ladders of 1 kb and ultra low resolution (ULR) DNA ladder; lane 3, fluorescent 6HB nanopores without cholesterol tags; lane 4, 6HB-3C fluorescent nanopores with three cholesterol tags; lane 5, 6HB-2C fluorescent nanopores with two cholesterol tags; lane 6, 6HB-1C fluorescent nanopores with one cholesterol tag. The position of the Kilobase pair length of dsDNA markers is indicated on the left of the gel. (b)Electrophoresis characterization (2% Agarose gel) of migration patterns of biotin modified nanopores in presence and absence of excess Streptavidin. Lanes 1 and 2, DNA ladders; lane 3, 6HB-2B nanopores without cholesterol modifications; lane 4, 6HB-2C nanopores without biotin modifications; lane 5, 6HB-2B-2C modifications; lane 6, 6HB-2B modifications with excess streptavidin (1x: 20x); lane 7, 6HB-2C modifications with excess streptavidin (1x: 20x); lane 8, 6HB-2C modifications with excess streptavidin (1x: 20x). The position of the base pair length of dsDNA markers are indicated in blue on the migration bands. (c) Dynamic light scattering trace of 6HB nanopores (blue line) without any cholesterol tags and 6HB-3C nanopores (orange line) containing three cholesterol tags.

**Supplementary Fig. 4.**
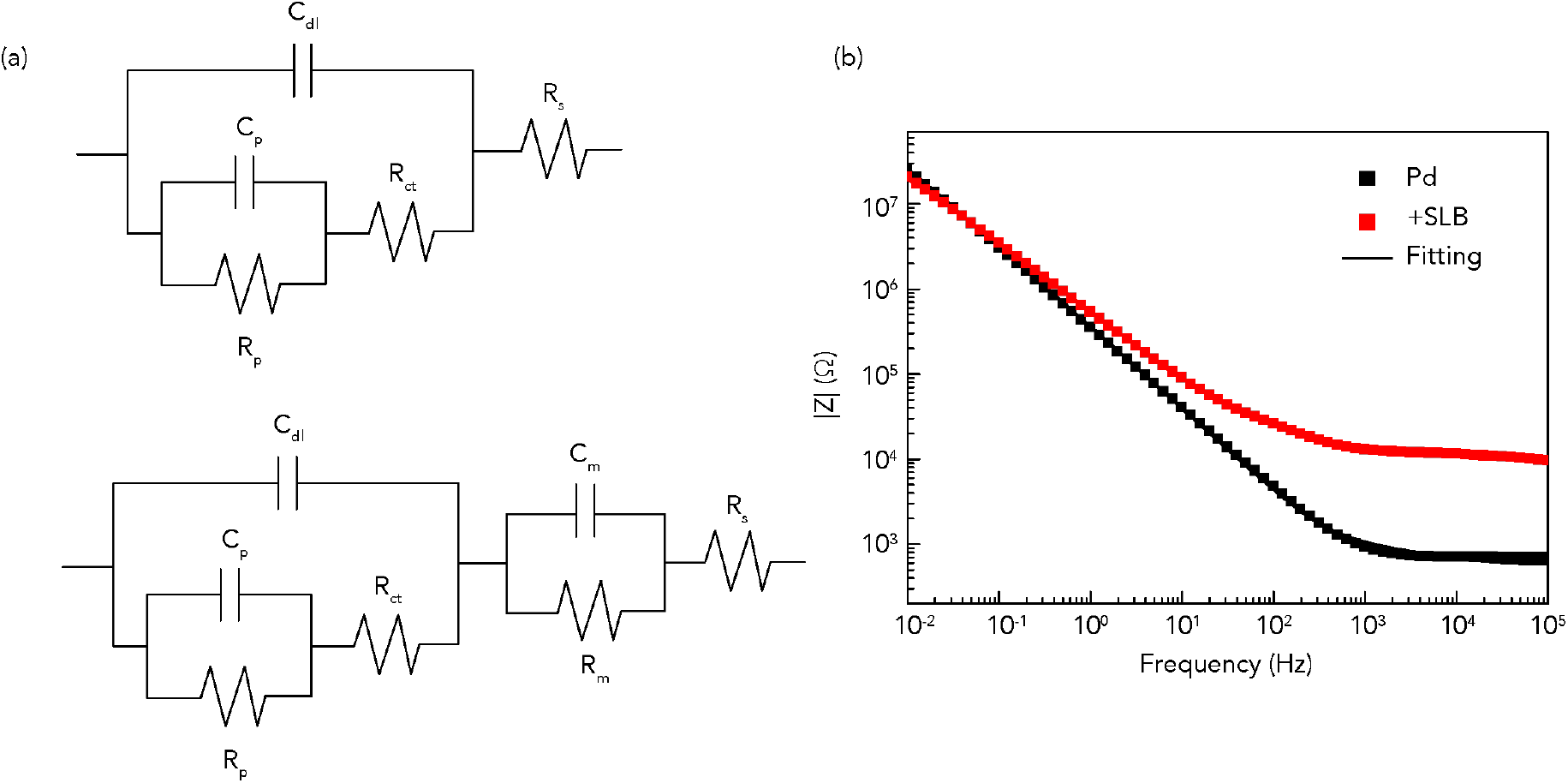
EIS measurement of bioprotonic device and lipid bilayer. (a) Equivalent circuit schematic for fit experimental data. (Top) Bioprotonic device, (Bottom) Bioprotonic device with SLB or SLB with DNA nanopore. The electrolyte solution resistance, R_s_, in series with membrane capacitance, C_m_, membrane resistance, R_m_, double layer capacitance, C_dl_, charge transfer resistance, R_ct_, adsorption resistance R_p_, and adsorption capacitance. (b) Bode plot (Black: Pd and Red: SLB).

**Supplementary Fig. 5.**
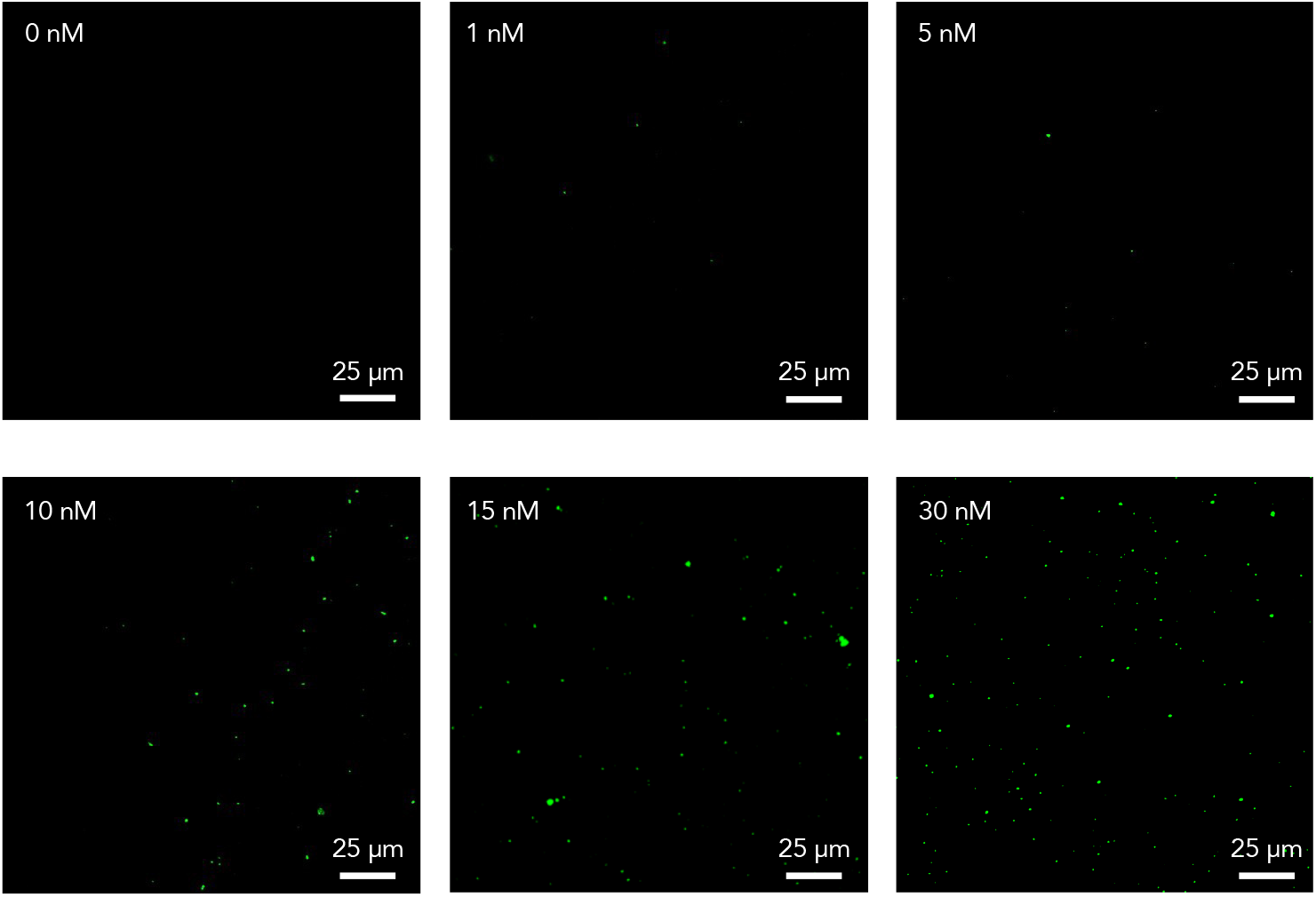
Fluorescence images of DNA nanopores at different concentrations.

**Supplementary Fig. 6.**
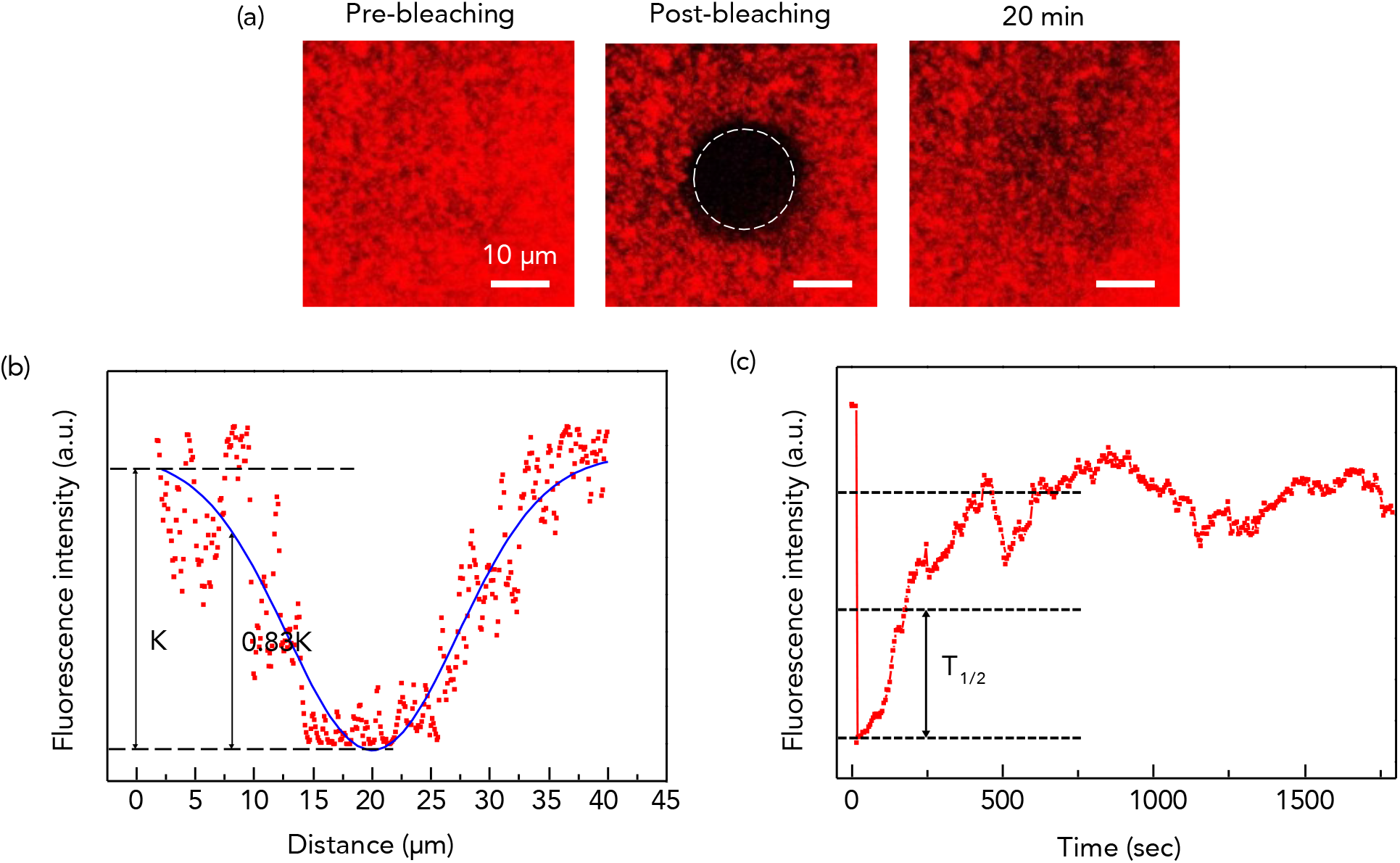
Characterization of lipid bilayer formation on Pd by FRAP. (a) Fluorescence intensity recovery after photobleaching is shown at t = pre, 0, and 20 min (from left to right) (b) Determination of effective bleaching spot. The Gaussian amplitude function was used to extract the effective bleaching spot radius. (c) The normalized fluorescence intensity of FRAP recovery curve.

**Table S1.**
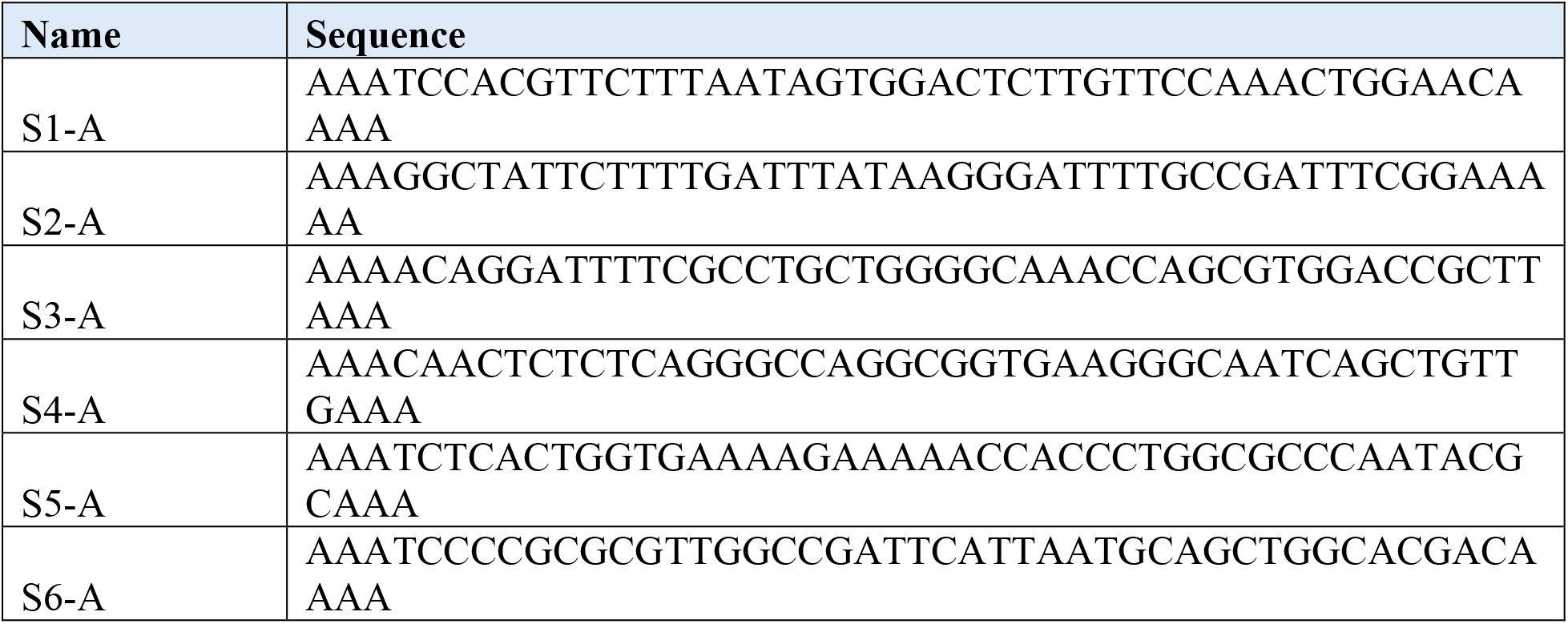
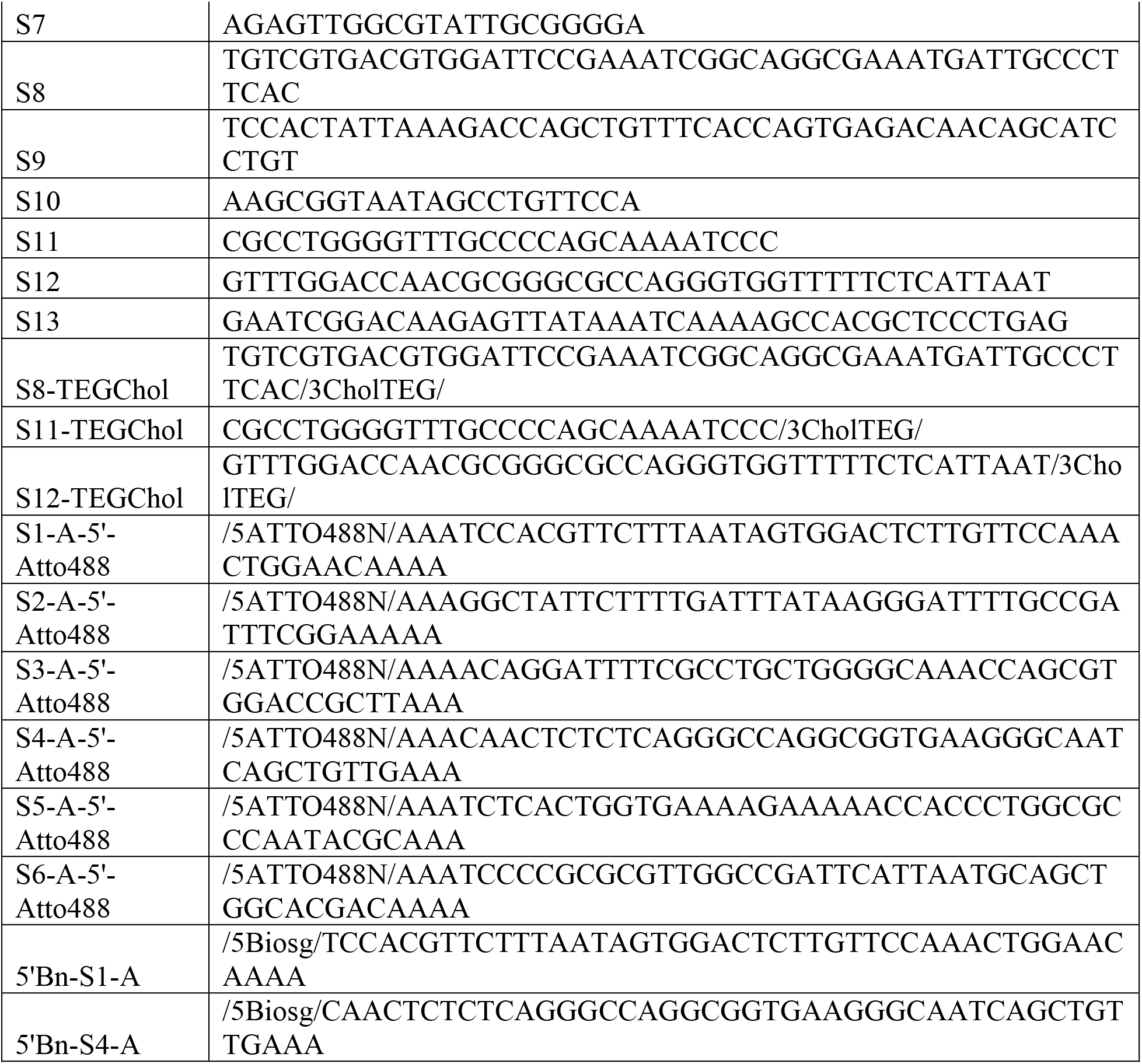
DNA oligo sequences.

## Acknowledgments

This work was supported by the National Science Foundation (NSF 20-518). We acknowledge using the Hyak supercomputer system at the University of Washington and Busra Demir acknowledges a TUBITAK 2214-A International Doctoral Research Fellowship.

## Author contributions

L.L., S.M. and Y.P. contributed equally to this work. A.G., M.R., and M.P.A conceived the biotic-abiotic interface concept. S.M. and L.L. designed the experiments. S.M. designed and characterized the DNA nanopores. L.L. and J.D. built the integrated electronic devices. L.L. and Y.P. conducted and analyzed the electronic and optical measurements. L.L., Y.P. and S.M. analyzed the data and formulated the analytical model with M.R. B.D. developed and analyzed the nanopore-bilayer simulation models. A.G. and M.R. supervised the experiments. E.E.O and M.P.A supervised the nanopore simulations. L.L., S.M., Y.P., B.D., and MR wrote the manuscript. M.R. and A.G. edited the manuscript and all authors read the manuscript.

## Competing interests

The authors declare no competing interests.

## Notes

### Competing Interest Statement

The authors have declared no competing interest.

