## Supplemental figure 1 will be used for the link to the file on the preprint site for "DNA nanopores as artificial membrane channels for origami-based bioelectronics"

**Merging electronics and biology with DNA based proton conducting membrane channels**

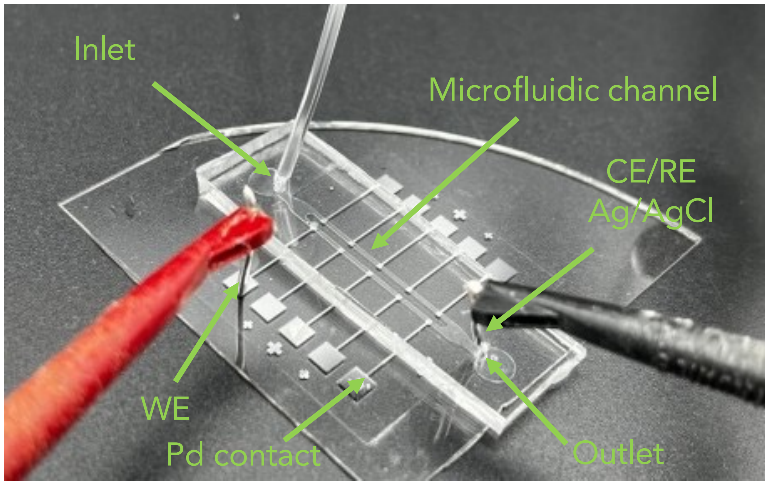


**Supplementary Fig. 1 Optical image of the bioprotonic device.**

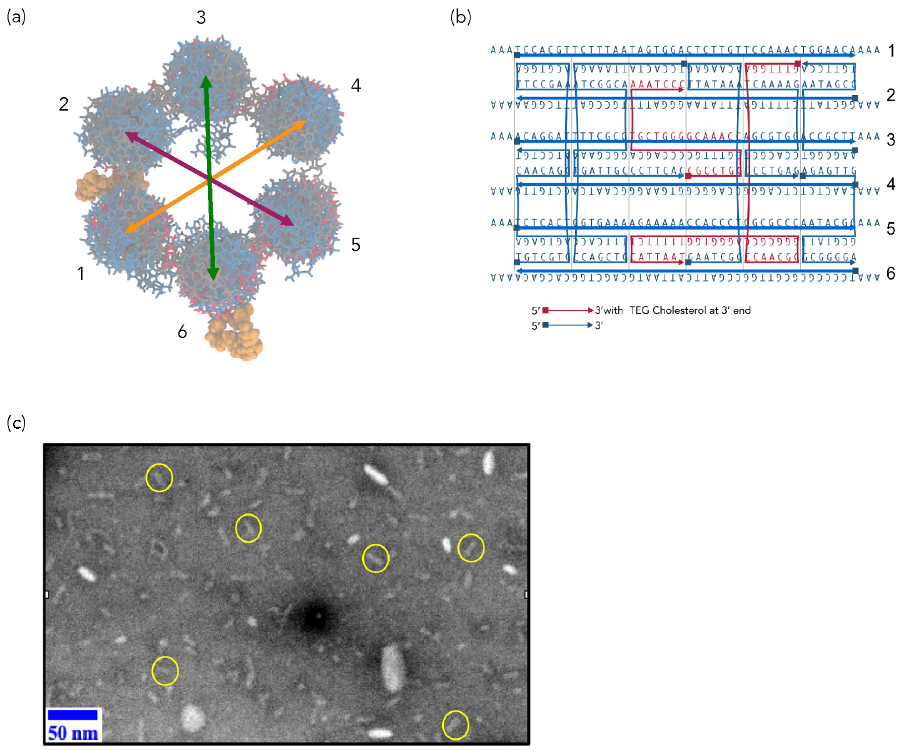


**Supplementary Fig. 2 6HB-2C nanopore design, conformation and simulations.** (a) Simulation of cholesterol moieties on helices 2 and 6 from an axial view with respect to the six helix bundles. (b) Sequence design and strand crossover details. Red strands indicate those oligos that have been modified at the 3' end with Tri-ethylene Glycol (TEG) cholesterol moieties. Blues strands indicate oligos without any modifications. Squares indicate the 5' end while triangles indicate the 3' end of DNA. (c) Negatively stained TEM micrograph of the 6HB nano-barrels. Yellow circles show the nanopores in a flat orientation.

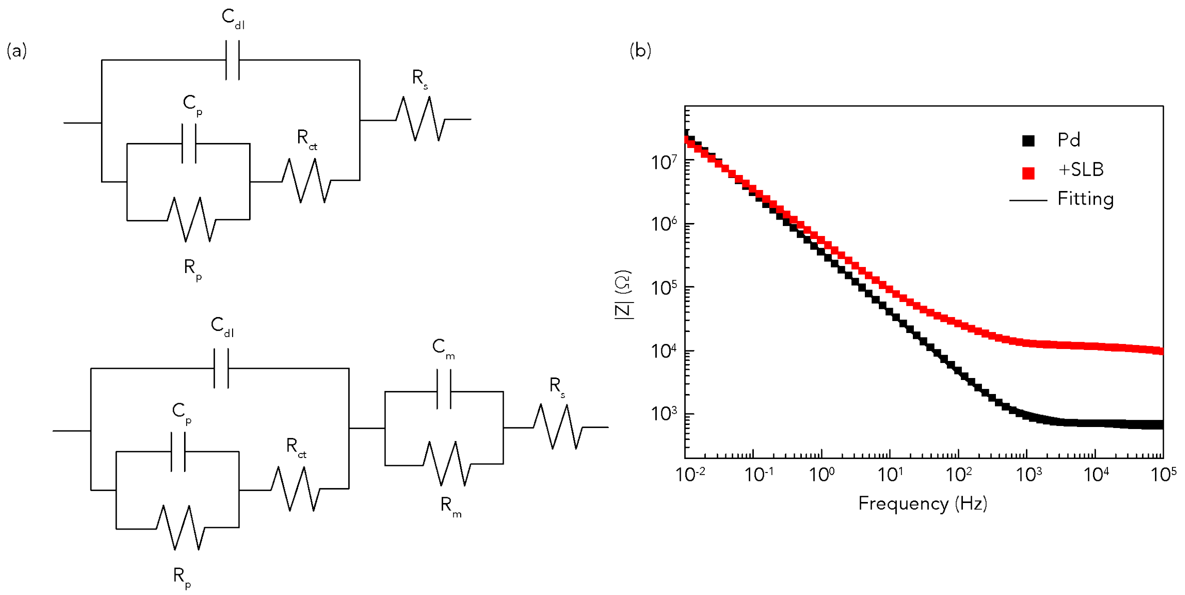


**Supplementary Fig. 3 EIS measurement of bioprotonic device and lipid bilayer.** (a) Equivalent circuit schematic for fit experimental data. (Top) Bioprotonic device, (Bottom) Bioprotonic device with SLB or SLB with DNA nanopore. The electrolyte solution resistance, $R_s$, in series with membrane capacitance, $C_m$, membrane resistance, $R_m$, double layer capacitance, $C_{dl}$, charge transfer resistance, $R_{ct}$, adsorption resistance $R_p$, and adsorption capacitance. (b) Bode plot (Black: Pd and Red: SLB)

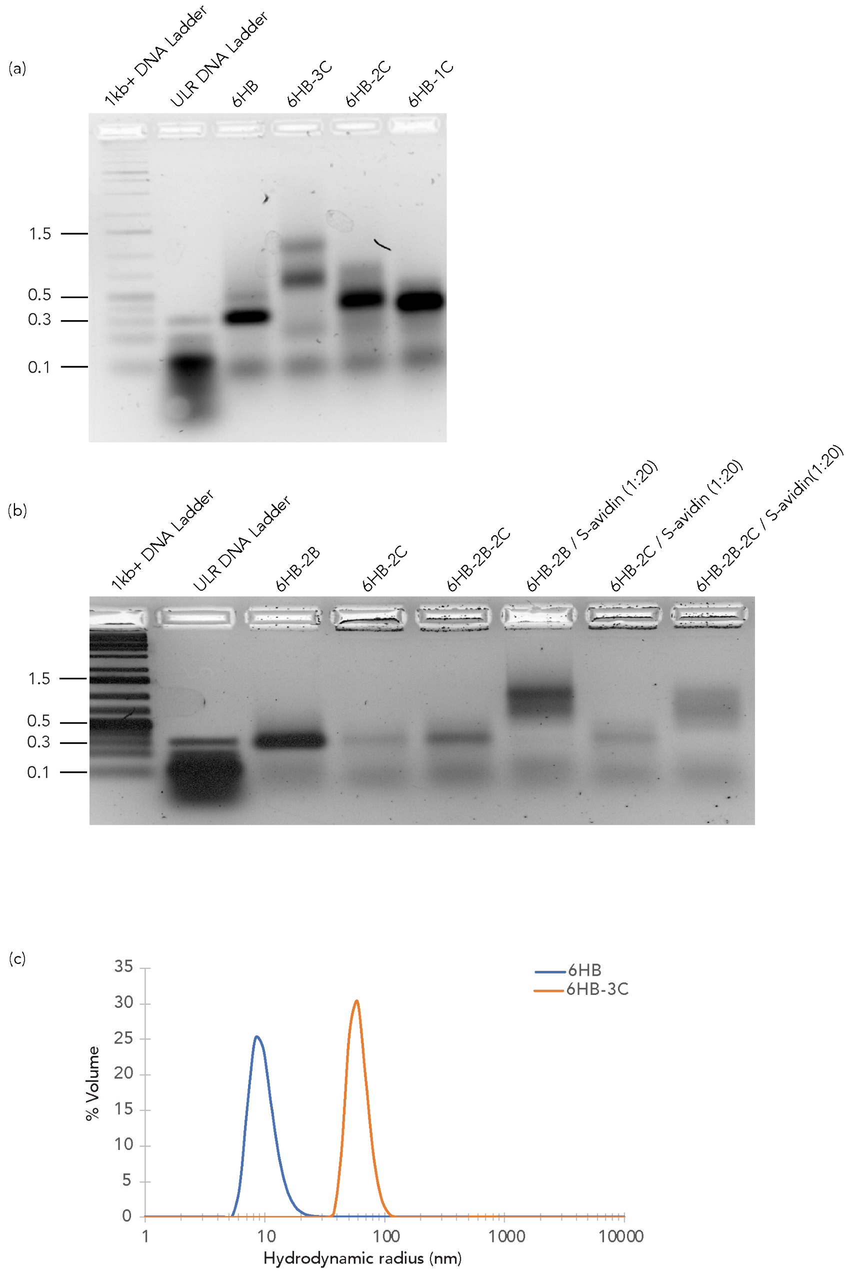


**Supplementary Fig. 4 Design and verification of DNA nanopores with cholesterol handles.** (a) Electrophoresis characterization (2% Agarose gel) of the nanopores with different number of cholesterol tags around the midsection of the nano-barrel. Lane 1 and 2, DNA ladders; lane 3, fluorescent 6HB nanopores without cholesterol tags; lane 4, 6HB-3C fluorescent nanopores with three cholesterol tags; lane 5, 6HB-2C fluorescent nanopores with two cholesterol tags; lane 6, 6HB-1C fluorescent nanopores with one cholesterol tag. The position of the Kilobase pair length of dsDNA markers is indicated on the left of the gel. (b) Electrophoresis characterization (2%

Agarose gel) of migration patterns of biotin modified nanopores in presence and absence of excess Streptavidin. Lanes 1 and 2, DNA ladders; lane 3, 6HB-2B nanopores without cholesterol modifications; lane 4, 6HB-2C nanopores without biotin modifications; lane 5, 6HB-2B-2C modifications; lane 6, 6HB-2B modifications with excess streptavidin (1x: 20x); lane 7, 6HB-2C modifications with excess streptavidin (1x: 20x); lane 8, 6HB-2C modifications with excess streptavidin (1x: 20x). The position of the base pair length of dsDNA markers are indicated in blue on the migration bands. (c) Dynamic light scattering trace of 6HB nanopores (blue line) without any cholesterol tags and 6HB-3C nanopores (orange line) containing three cholesterol tags.

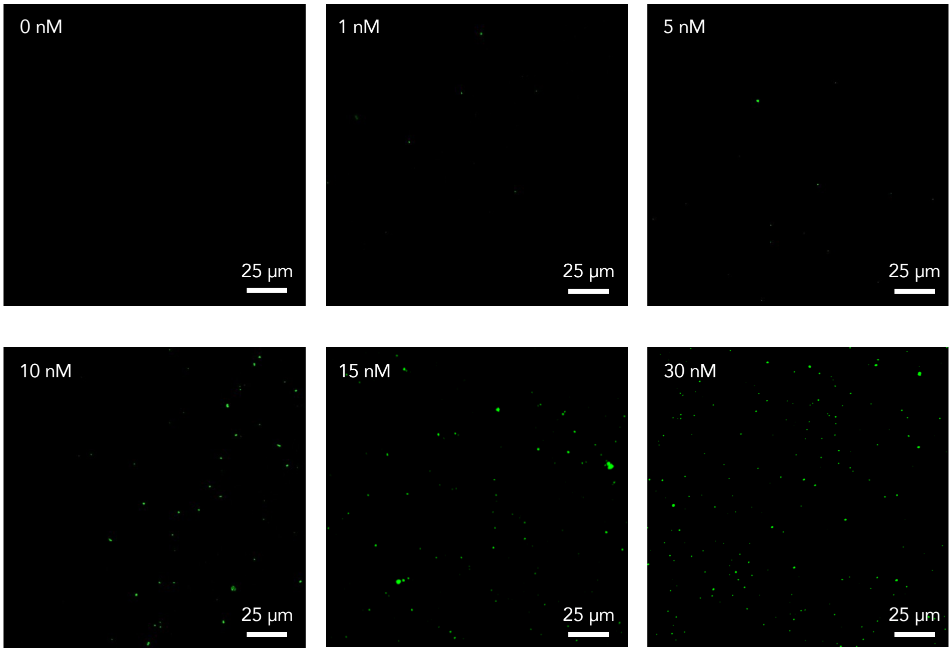


**Supplementary Fig. 5 Fluorescence images of DNA nanopores at different concentrations.**

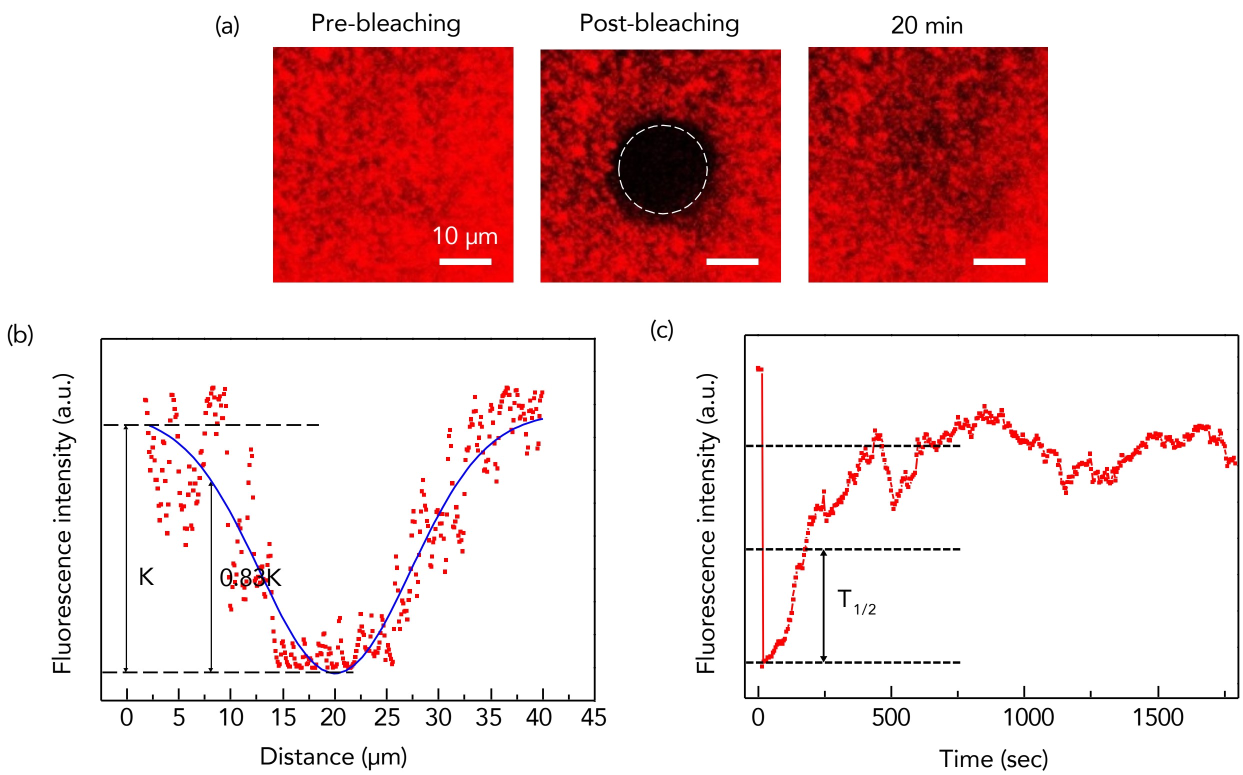


**Supplementary Fig. 6 Characterization of lipid bilayer formation on Pd by FRAP.** (a) Fluorescence intensity recovery after photobleaching is shown at t = pre, 0, and 20 min (from left to right) (b) Determination of effective bleaching spot. The Gaussian amplitude function was used to extract the effective bleaching spot radius. (c) The normalized fluorescence intensity of FRAP recovery curve.

**Table S1. DNA oligo sequences**

| Name | Sequence |
|---|---|
| S1-A | AAATCCACGTTCTTTAATAGTGGACTCTTGTTCCAAACTGGAACAAAA |
| S2-A | AAAGGCTATTCTTTTGATTTATAAGGGATTTTGCCGATTTCGGAAAAA |
| S3-A | AAAACAGGATTTTCGCCTGCTGGGGCAAACCAGCGTGGACCGCTTAAA |
| S4-A | AAACAACTCTCTCAGGGCCAGGCGGTGAAGGGCAATCAGCTGTTGAAA |
| S5-A | AAATCTCACTGGTGAAAAGAAAAACCACCCTGGCGCCCAATACGCAAA |
| S6-A | AAATCCCCGCGCGTTGGCCGATTCATTAATGCAGCTGGCACGACAAAA |

| | |
|---|---|
| S7 | AGAGTTGGCGTATTGCGGGGA |
| S8 | TGTCGTGACGTGGATTCCGAAATCGGCAGGCGAAATGATTGCCCTTCAC |
| S9 | TCCACTATTAAAGACCAGCTGTTTCACCAGTGAGACAACAGCATCCTGT |
| S10 | AAGCGGTAATAGCCTGTTCCA |
| S11 | CGCCTGGGGTTTGCCCCAGCAAAATCCC |
| S12 | GTTTGGACCAACGCGGGCGCCAGGGTGGTTTTTCTCATTAAT |
| S13 | GAATCGGACAAGAGTTATAAATCAAAAGCCACGCTCCCTGAG |
| S8-TEGChol | TGTCGTGACGTGGATTCCGAAATCGGCAGGCGAAATGATTGCCCTTCAC/3CholTEG/ |
| S11-TEGChol | CGCCTGGGGTTTGCCCCAGCAAAATCCC/3CholTEG/ |
| S12-TEGChol | GTTTGGACCAACGCGGGCGCCAGGGTGGTTTTTCTCATTAAT/3CholTEG/ |
| S1-A-5'-Atto488 | /5ATTO488N/AAATCCACGTTCTTTAATAGTGGACTCTTGTTCCAAACTGGAACAAAA |
| S2-A-5'-Atto488 | /5ATTO488N/AAAGGCTATTCTTTTGATTTATAAGGGATTTTGCCGATTTCGGAAAAA |
| S3-A-5'-Atto488 | /5ATTO488N/AAAACAGGATTTTCGCCTGCTGGGGCAAACCAGCGTGGACCGCTTAAA |
| S4-A-5'-Atto488 | /5ATTO488N/AAACAACTCTCTCAGGGCCAGGCGGTGAAGGGCAATCAGCTGTTGAAA |
| S5-A-5'-Atto488 | /5ATTO488N/AAATCTCACTGGTGAAAAGAAAAACCACCCTGGCGCCCAATACGCAAA |
| S6-A-5'-Atto488 | /5ATTO488N/AAATCCCCGCGCGTTGGCCGATTCATTAATGCAGCTGGCACGACAAAA |
| 5'Bn-S1-A | /5Biosg/TCCACGTTCTTTAATAGTGGACTCTTGTTCCAAACTGGAACAAAA |
| 5'Bn-S4-A | /5Biosg/CAACTCTCTCAGGGCCAGGCGGTGAAGGGCAATCAGCTGTTGAAA |
